# Sub-anesthetic ketamine administration decreases deviance detection responses at the cellular, populational and mesoscale levels

**DOI:** 10.1101/2025.09.05.674559

**Authors:** Maria Isabel Carreño-Muñoz, Alessandra Ciancone Chama, Pegah Chehrazi, Bidisha Chattopadhyaya, Graziella Di Cristo

**Author notes:** Corresponding authors: Maria Isabel Carreño-Muñoz, Department of Neurosciences, Université de Montréal, CHU Sainte-Justine Azrieli Research Center, 3175, Côte-Sainte-Catherine, Montréal, QC H3T 1C5, Canada., Graziella Di Cristo, Department of Neurosciences, Université de Montréal, CHU Sainte-Justine Azrieli Research Center, 3175, Côte-Sainte-Catherine, Montréal, QC H3T 1C5, Canada. Equal contribution (Maria Isabel Carreño-Muñoz, Alessandra Ciancone Chama).

## Abstract

In the neocortex, neuronal processing of sensory events is significantly influenced by their predictability. A common example is the suppression of responses to repetitive stimuli in sensory cortices, a phenomenon known as habituation. Within a sensory information stream, whenever a novel stimulus deviates from expectations, enhanced brain responses are observed. Mismatch negativity (MMN), the electroencephalographic waveform reflecting rule violations, is a well-established biomarker for auditory deviant detection. MMN has been shown to depend on intact NMDA receptor signaling across species; nevertheless, the underlying mechanisms at the neuronal and mesoscale levels are still not fully understood. Using multi-electrode array recordings in awake mice, we identified a specific biphasic spiking response in a subpopulation of primary auditory cortex (A1) neurons elicited by deviant, but not standard, sounds, wherein the second peak is abolished by acute sub-anesthetic injection of ketamine, a partial non-competitive NMDA receptor antagonist. We further showed that the posterior parietal cortex (PPC), a critical hub for multisensory integration and sensorimotor coordination, responds to deviant, but not repetitive, sounds, and this response is dependent upon intact NMDA receptor-mediated signaling. Finally, to explore the effects of ketamine on inter-cortical communication following deviance detection, we performed Weighted Phase Lag Index (wPLI) analyses during the presentation of deviant and standard sounds. This analysis showed a functional connectivity between A1 and PPC following deviant detection, which is impaired by ketamine administration. Altogether, our findings provide novel insights into the NMDA receptor-dependent mechanisms underlying the processing of novelty in auditory stimuli.

## INTRODUCTION

Animal survival critically depends on the continuous updating of sensory information and its rapid processing to enable appropriate behaviors. To interact efficiently with ever-changing environments, mammalian brains must continuously align incoming sensory information with predictions based on current actions or intentions. Consequently, the brain has evolved various strategies to minimize computational demands while still efficiently navigating through a constant flow of information. One such strategy is the ability to suppress responses to predictable stimuli — a phenomenon referred to here as habituation, or stimulus-specific adaptation — while remaining alert to unexpected changes – referred to as deviance detection. In sensory cortices, neurons gradually reduce their firing to repetitive sounds but preserve robust responses to rare, deviant stimuli (Harms *et al*., 2016). This form of neuronal adaptation also manifests as changes in evoked related potentials (ERPs), which are time-locked brain responses that represent changes at the population level in recorded local field potentials (LFP). As it is the case for neuronal firing, repetitive sounds also elicit a progressive reduction in ERP amplitude (Rosburg & Mager, 2021), while unexpected sounds trigger a specific wave shape response known as mismatch negativity (MMN)(Naatanen *et al*., 2007), reflecting the brain’s detection of deviations from auditory regularities. Together, auditory habituation and MMN both at the cellular and population level, provide a hierarchical framework for understanding how local neuronal response and large-scale network dynamics contribute to efficient sensory monitoring and predictive coding in the brain.

At the inter-cortical (mesoscale) level, recent studies have proposed that both functional and anatomical connectivity between distinct brain regions are essential for processing standard and deviant auditory stimuli. While several studies have emphasized the role of frontal cortex in top-down modulation of sound predictability (Hockley *et al*., 2025), emerging evidence also highlights the potential involvement of the posterior parietal cortex (PPC), a higher-order associative region involved in both multisensory integration and sensorimotor coordination (Andersen & Buneo, 2002; Van Derveer *et al*., 2023). Studies in rodents and humans have shown that the PPC exhibits robust responses to unexpected stimuli in the auditory domain (Justen & Herbert, 2018; Raltschev *et al*., 2025). Further, at the cellular level, PPC neurons can discriminate the presence or absence of expected auditory stimuli, in awake mice (Mohan *et al*., 2018). However, it is still unknown how the PPC communicates with other cortical areas to contribute, at the mesoscale level, to the processing of rule violations.

Although the synaptic mechanisms underlying rule-violation processing are not completely understood yet, most studies propose N-methyl-D-aspartate (NMDA) receptor activity as a central mechanism (Light & Naatanen, 2013; Rosburg & Kreitschmann-Andermahr, 2016). For example, studies using sub-anesthetic doses of ketamine, a partial non-competitive antagonist of NMDA receptors, reliably reported reductions in MMN amplitude and auditory habituation in humans (Umbricht *et al*., 2000; Kreitschmann-Andermahr *et al*., 2001; Rosburg & Kreitschmann-Andermahr, 2016). While the effects of ketamine administration on MMN and auditory habituation at the local population level are relatively well documented, the impact of ketamine at the cellular and mesoscale levels, particularly in relation to functional inter-cortical connectivity and network dynamics, remains largely unexplored.

In this study, we assessed how a sub-anesthetic dose of ketamine affects auditory deviant detection and habituation at the population level, through ERP-MMN waveforms and evoked oscillatory activity analysis, and at the neural level, by analyzing single-unit firing, in the primary auditory cortex (A1) of awake mice. Furthermore, we explored the functional connectivity between PPC and A1 to gain a deeper understanding of the broader network dynamics involved both in auditory deviant detection and habituation.

## MATERIAL AND METHODS

### Animals

Experimental procedures involving mice were approved by the “Comité Institutionnel de Bonnes Pratiques Animales en Recherche” (CIBPAR) of the Research Center of Sainte-Justine Hospital in accordance with the principles published by the Canadian Council on Animal Care. C57Bl/6 mice were implanted at postnatal day (P)70–90 and recorded between P80 and P200. All animals were housed with two to five mice/cage from weaning until surgery, after which they were singly housed for the duration of the experiment. Animals were kept on a 12h/12h light/dark cycle and provided with nesting material, food, and water *ad libitum*. All experiments were performed during the light period under constant mild luminosity (60 lx). We checked and noted the exact stage of the females’ estrous cycle after each recording session by histological inspection of the vaginal smear (McLean *et al*., 2012). To avoid variability and the concomitant effect of sex hormones, the two females included in our analysis were in the diestrus phase in all conditions (control, vehicle and ketamine).

All mice were recorded in three experimental conditions: naïve (no injection), following vehicle injection, and following ketamine injection. Intraperitoneal (IP) sub-anesthetic dose of ketamine (8 mg/Kg diluted in saline solution) or vehicle (saline solution) were injected 10 minutes before the start of the recording session. The administration of vehicle and ketamine was randomized, so that one group mice received the vehicle injection first and then the ketamine injection after, while a second group of mice received the ketamine injection first followed by the vehicle injection.

### Surgery

Cranial surgery was performed under deep anesthesia, induced by an intraperitoneal injection of ketamine/xylazine cocktail (ketamine 100 mg/Kg and xylazine 10 mg/Kg). Body temperature was maintained at 37°C using a feedback-controlled heating pad. Analgesics were given locally subcutaneously (Lidocaine 2%, Aspen Pharmacare, Canada) and an intraperitoneally dose of 30 mg/Kg (Buprenorphine, 0,3mg/ml, Vetergesic Multidose, Ceva) 5 min before surgery. Mice were head-fixed in a stereotaxic frame (Kopf Instruments, USA). During the entire surgery, eyes were protected and hydrated using Opticare eye lube (CLC Medica, Canada). The skull was exposed, cleaned, and treated with an activator (Metabond Quick, Parkell, USA) to increase bone porosity. The coordinates of the targeted areas (described below) were marked with a permanent marker before fixing the skin around the exposed skull with Vetbond tissue adhesive (3M, USA). Then a first layer of dental cement (Metabond Quick, Parkell, USA) was applied to cover the whole exposed skull except the marked coordinates. Craniotomies were then performed above the targeted areas.

In the first set of experiments, custom-made clusters of four to six 100 µm-diameter-insulated-tungsten wires (50 µm-diameter bared wires) were implanted in every targeted area. In each one of those clusters, wire tips were 50–150 µm apart. A custom-designed and 3D-printed microdrive-like scaffold held the three electrode clusters at fixed positions (Supplemental Figure 1A, B) as previously described in (Carreno-Munoz *et al*., 2022), thus allowing the simultaneous implantation of all three clusters and reducing surgery time. We used only recording electrodes showing impedances in a range between 100-400 KΩ. In a second set of experiments, four independently movable tetrodes targeted the primary auditory cortex. Each tetrode was fabricated out of four 10 μm-diameter tungsten wires (H-Formvar insulation with Butyral bond coat California Fine Wire Company, Grover Beach, CA) that were twisted and then heated to bind them into a single bundle. The tips of the tetrodes were then gold-plated to reduce the impedance to <400 kΩ. The use of tetrodes provides not only local field potential (LFP) recordings but also high-density spiking activity, allowing for the characterization of single-unit discharges. The 3D printed microdrives used were manufactured by Axona (Axona Ltd.; http://www.axona.com/).

A ground-and-reference wire was also gently introduced in the contralateral frontal lobe. The microdrive apparatus and the ground-and-reference wire were then daubed with dental acrylic (Ortho-Jet Powder and Jet liquid, Lang Dental, USA) to encase the electrode-microdrive assembly and anchor it to the skull. When implanting tetrodes, a drop of silicone elastomer (WPI, USA) was applied around the tetrodes before the acrylic to let tetrodes free for movement. For 1 week after surgery, animals were treated with Metacam and the skin around the implant was inspected daily.

### Coordinates

Electrode clusters were implanted at the following coordinates: primary auditory cortex (A1), −2.5 mm posterior to bregma, 3.9 mm lateral to the midline and 0.8 mm deep; primary visual cortex (V1), −4 mm posterior to bregma, 2 mm lateral to the midline, depth: 0.6 mm; posterior parietal cortex (PPC), −2 mm posterior to bregma,1.7 mm lateral to the midline, depth: 0.6 mm. Clusters placed in the V1 and PPC targeted both the supra and infragranular layers; longest wire from PPC cluster also reached superficial layers from the CA1 hippocampal region placed above. Clusters placed in the A1 only targeted layer 5. Data shown in this study are specifically from layer 5 A1 and PPC. To verify the electrode cluster locations, after the recording experiments, mice were perfused transcardially with saline followed by 4% paraformaldehyde in phosphate buffer (PB, 0.1 M, pH 7.2). Brains were post-fixed with 4% paraformaldehyde overnight and the implanted electrodes were carefully removed. Subsequently the brains were transferred to a 30% sucrose solution in PBS for 48 h. The tissue was then embedded in OCT compound (Sakura Finetek, Torrance, CA, USA) prior to cryosectioning. Coronal sections (40 µm thick) were obtained using a cryostat (Leica Microsystems VT100). Brain sections were then labeled with DAPI (1:1000 in PBS) for 25 min at room temperature to visualize cell nuclei. After rinsing three times with PBS, the slices were mounted in Vectashield mounting medium (Vector). Whole-brain slices were imaged on a wide-field fluorescent microscope (Leica DMi8) using 10X (0.3 NA) and 20X (0.8 NA) objectives (Supplemental Figure 1C, D). For each slice, 10X images of the entire slice were acquired as a series of overlapping tiles and stitched automatically to generate a single high-resolution image. Higher-magnification images were acquired at the auditory and posterior parietal cortex coordinates to confirm electrode location.

### LFP data acquisition

LFP recordings were performed using an open-ephys GUI platform (https://open-ephys.org/). Data was acquired at 20 KHz with a custom-made headstage using Intan Technologies RHD Electrophysiology Amplifier chip (https://intantech.com/products_RHD2000.html). Custom designed auditory and visual trigger generator devices sent triggers directly to the recording system, assuring the precise timing of stimuli presentation. Starting 1 week after surgery, animals were habituated daily for 4 days to an open field (15–20 min each time), before starting recordings for auditory stimulation protocols from Day 5. We used a dimly illuminated open field environment (45 × 35 cm), surrounded by 60 cm-high transparent walls, equipped with video monitoring. The walls were sprayed and wiped clean with Peroxigard (Virox Technologies) 15 min before the introduction of each animal. Two speakers were placed ∼10 cm from the walls of the open field. Baseline activity was recorded at the beginning of each session for a period of 10 min, while mice were freely exploring the open field. Data acquisition during auditory stimulations was conducted for 20–30 min, a total of three to six times, on alternate days.

### Auditory stimulation protocols

All stimuli were presented at an intensity of 70-75 dB. The oddball paradigm comprised a sequence of four to nine repetitive (standard) sounds at 5 kHz, followed by a deviant sound at 10 kHz. Six different sequences (containing 4/1, 5/1, 6/1, 7/1, 8/1 or 9/1 standard/deviant sounds) were presented 120 times in a randomized order. The total number of stimulations was 450 and each sound was presented 60 (S1, S2, S3, S4 and D), 50 (S5), 40 (S6), 30 (S7), 20 (S8) or 10 (S9) times to all animals. Sounds lasted 70ms and were presented with a 1s interstimulus interval. The full protocol lasted 10 min. For the reversed oddball protocol, the stimulus probabilities were swapped (standard sound was 10 kHz and deviant sound was 5kHz), but the sequences were organized similarly.

### LFP preprocessing

LFP signals were down-sampled to 2000 Hz and a bandpass filter (0.5–150 Hz) was applied. A notch filter (59.5–60.5 Hz) was also applied to remove residual 60 Hz noise contamination. Data was then segmented into 1400ms (500ms pre- and 900ms post-stimulus onset) periods. Trials with an intra-channel average higher than four times the total trial standard deviation (SD) were tagged, visually inspected and removed. Only sessions containing more than 50 clean trials were kept for further analyses.

### Data analysis

None of the recorded animals were excluded from the final analysis. Signal analysis and quantification were performed using custom MATLAB (The Mathworks Inc.) codes, available upon request. Animal behavior was analyzed using DeepLabCut (Mathis *et al*., 2018). Body center was extracted and used to measure mice locomotor activity.

#### Evoked Related Potentials

The LFP signal was baseline corrected to the mean voltage at 100ms prior to stimulus onset and averaged over trials. The evoked related potentials (ERPs) components, baseline-to-peak N1, and P1, were analyzed from the grand-average. The N1 amplitude was automatically detected by subtracting the minimum voltage (negative peak) within a 10-60ms time window after stimulus onset to the average baseline value. The P1 amplitude was calculated by measuring the maximum voltage (positive peak) within an 80-150ms time window after stimulus onset to the average baseline value. Analyses were carried out using the Fieldtrip toolbox (Oostenveld *et al*., 2011). To quantify habituation from ERPs, we calculated the logarithmic ratio between the N1 amplitudes of the standard before the deviant (SBD) and the first standard (S1) sounds by dividing S1 by SBD. These ratios were then converted to a logarithmic base 10 scale and multiplied by 10, using the following formula:

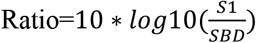

Mismatch negativity (MMN) trace was calculated by subtracting the LFP trace of the standard sound before the deviant (SBD) from the LFP trace of the deviant sound (D). From the pattern obtained from each subject’s ERPs we calculated the average value between 77ms and 84ms following stimulus presentation to obtain the MMN response, which corresponds to the most clearly detectable peak in the MMN waveform.

#### Power spectral density

Baseline power spectral density (PSD) analysis, was done during periods of immobility within the first 10 minutes of the recording session in the absence of any auditory stimuli. Immobility periods were defined by establishing a velocity threshold of the animal body center at <1 cm/s for at least 2 seconds. To estimate the power spectrum using Fast Fourier Transform (FFT), we used the Welsh method implemented in MATLAB. The estimated power spectrum was normalized by the total power of the signal and plotted using a logarithmic scale for visualization purposes.

Evoked PSD analysis was performed on the 300ms following stimulation for MMN and on the 250ms following stimulation for habituation. These were baseline corrected using the same length periods for corresponding baselines through the formula:

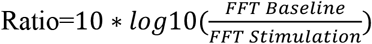

To adopt an approach comparable to the MMN waveform and facilitate the comparison of habituation and deviant detection between experimental groups, we calculated the difference (Δ) in evoked PSD by subtracting the corresponding sound conditions: D − SBD (deviant detection) and S1 – SBD (habituation).

#### Spiking activity analysis

Offline processing of electrophysiological data began with the extraction of spikes from the high-pass filtered signal (800–5000 Hz) of recordings sampled at 20 kHz. Action potentials exceeding 5 standard deviations above the mean baseline were detected. Spike waveforms were then characterized by feature extraction, and putative neuronal units were identified using principal component analysis followed by automatic clustering using klustakwik (Harris *et al*., 2000) (http://klustakwik.sourceforge.net/). Identified clusters were manually refined with a graphical cluster-cutting program (Csicsvari *et al*., 1998). Only units exhibiting a clear refractory period (<2ms) in their auto-correlograms and well-isolated cluster boundaries were included in subsequent analyses.

Peri-stimulus time histograms (PSTH) and spike density function (SDF) were performed using the mlib toolbox by Maik Stüttgen (https://de.mathworks.com/matlabcentral/fileexchange/37339-mlib-toolbox-for-analyzing-spike-data). PSTHs were calculated using a bin size of 5 ms. SDF were calculated by applying a gaussian distribution over the PSTH using a kernel of 5 ms width. Each neuron SDF was baseline corrected by subtracting SDF values during 50ms before the stimulus presentation from the entire SDF. Using this analysis, we identified 3 types of neurons, based on their firing behavior following the stimulus presentation: neurons that respond by increasing their firing (responding-excited), neurons that respond by decreasing their firing (responding-inhibited) and neurons whose firing does not change (non-responding neurons).

To facilitate the comparison of habituation and deviant detection between experimental groups, we calculated the difference (Δ) SDF by subtracting individual neurons SDF following deviant sound from SDF following standard sounds.

#### Connectivity analysis

Weighted Phase Lag Index (wPLI) analysis between A1 and PPC was also performed using the fieldtrip toolbox (Oostenveld *et al*., 2011). This analysis was performed during the MMN response period (80-320ms after the stimulus presentation) to avoid the first abrupt N1 peak and therefore the non-stationarity of the LFP signal. The wPLI was calculated using a Fourier-based non-parametric spectral factorization approach, employing a multi-taper frequency transformation with DPSS tapers of order 5 and a padding factor of 3. As before, we calculated the difference (Δ) in wPLI spectrograms by subtracting the spectrogram calculated during the periods following deviant sound (D) stimulation from those following standard sound before the deviant sound (SBD) stimulations.

#### Statistical Analysis

LFP were analyzed by a non-parametric repeated-measure test, the Wilcoxon Signed Rank Test. This test was chosen to avoid assuming normal distribution of the data. Spiking activity was analyzed by LMM, modelling animal as a random effect and treatment as fixed effect. We used this statistical analysis because we considered the number of mice as independent replicates and the number of cells in each mouse as repeated measures (Yu *et al*., 2022). Statistical analysis was performed using IBM SPSS V29.0.0 for LMM analysis. Results were considered significant when p<0.05. All data represent mean ± SEM. All data are available upon request.

## RESULTS

### Sub-anesthetic ketamine administration significantly reduces MMN response and auditory habituation

To validate previous findings reporting a reduction of MMN responses in humans (Rosburg & Kreitschmann-Andermahr, 2016), primates (Gil-da-Costa *et al*., 2013) and rodents (Schuelert *et al*., 2018), we first assessed the effect of an acute sub-anesthetic dose of ketamine (8 mg/Kg diluted in saline solution) or vehicle (saline solution) on auditory MMN response and habituation in freely moving mice. The dose of ketamine selected was based on previous preclinical studies showing its rapid effects on cortical activity (Picard *et al*., 2019). Six animals implanted with intracranial electrodes and familiarized with an open field arena were recorded before and 10 mins after ketamine or vehicle IP (intraperitoneal) injection, during an auditory oddball paradigm consisting of a series of repetitive (standard) sounds randomly interrupted by deviant sounds (Figure 1A). We calculated MMN by subtracting the LFP trace of the standard sound before the deviant (SBD) from the LFP trace of the deviant sound (D). The positive deflection in this waveform, peaking at approximately 100 ms after stimulus onset, is known as the MMN response, and it has been proposed to reflect the brain’s automatic detection of rule-violations of auditory regularities. Our results show that ketamine, but not vehicle, administration significantly reduced the amplitude of MMN response (Figure 1C, D). This data is consistent with previously reported observations in humans (Rosburg & Kreitschmann-Andermahr, 2016), primates (Gil-da-Costa *et al*., 2013) and rodents (Schuelert *et al*., 2018), thus validating our model. The effect of ketamine was not dependent of the stimulus frequency, since ketamine, but not vehicle, administration significantly reduced the amplitude of MMN response when using a reversal oddball sequence, too (Supplemental Figure 2).

Acute ketamine or vehicle injections did not significantly affect auditory habituation to standard sounds (Figure 1F), measured as the ratio between the N1 amplitude of the standard before the deviant (SDB) and the first standard sounds (S1). Since higher doses of ketamine administration can induce hyperlocomotion (de la Salle *et al*., 2022) and previous studies showed that auditory processing is modulated by motor activity (Schneider *et al*., 2014; Bigelow *et al*., 2019), we measured the total distance traveled by each mouse during the whole recording session. We found no effect of ketamine or vehicle injection on mouse locomotion (Figure 1G), thus ruling out an effect of altered motor activity on auditory processing.

### Sub-anesthetic ketamine administration significantly reduces the late component of spiking activity response

To further understand the effect of ketamine on MMN responses at the cellular level, we analyzed concomitant neuronal activity during oddball protocol stimulation after vehicle or ketamine injection. To do so, we identified spiking activity of individual neurons and evaluated their firing rates by calculating their peri-stimulus time histograms (PSTH, Figure 2A, lower panels) and spike density functions (Figure 2B). Our results show three different populations, based on their firing behavior following the deviant sound onset: neurons that respond by increasing their firing (responding-excited, Figure 2A-D), neurons that respond by decreasing their firing (responding-inhibited, Figure 2E-H) and neurons whose firing does not change (non-responding neurons, therefore not analyzed further). In naïve mice, deviant, but not standard sounds induced a biphasic response in the spiking activity of A1 neurons that were excited by the sounds (Figure 2B). This response was characterized by a fast and early burst of activity peaking at 25ms after stimulus onset, followed by a slow and late response at 100-200 ms after stimulus onset, sharing common temporal dynamics with the MMN response. This observation is in line with previous reports (Chen *et al*., 2015), suggesting that the second late component of spiking activity may code for novelty in A1 neurons (Ulanovsky *et al*., 2003) as well as those in other regions such as in the substantia nigra (Mikell *et al*., 2014) and PFC (Camalier *et al*., 2019). To facilitate comparison between experimental groups, we calculated the difference (Δ) in spike density functions (SDF) by subtracting the standard before the deviant sound condition (SBD) from the deviant sound condition (D). We refer to this subtractive measure as Δ SDF (Figure 2C, 2G). Analysis of Δ SDF showed that ketamine, but not vehicle, administration completely abolished this novelty-related late component of the spiking activity response (Figure 2D), therefore specifically affecting the saliency of the novel stimuli but not its deviant nature within the sequence of sounds.

**Figure 1.**
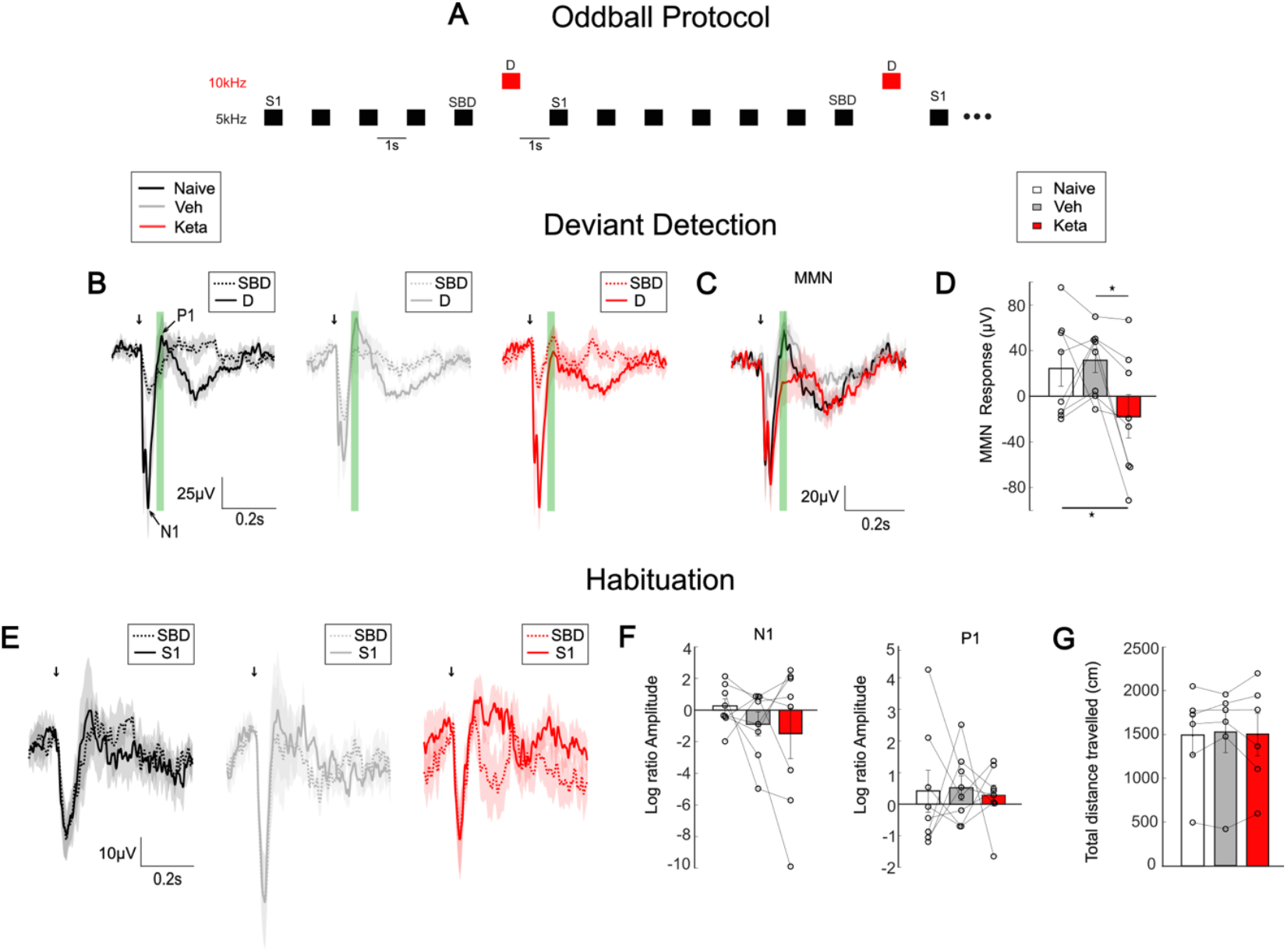
Subanesthetic ketamine alters MMN responses during deviant detection and stimulus specific adaption. (**A**) Schematic representation of the oddball protocol with black squares representing 5kHz pure tones (standard sounds) and the red squares 10kHz pure tones (deviant sounds). (**B**) Mean evoked related potentials (ERP) traces of standard before the deviant (SBD, dotted) and deviant (D, solid) sounds for naive (left), vehicle injected (middle), and ketamine injected (right) mice. Arrows indicate the start time of stimulation. N1 is identified as the first negative peak and P1 as the first positive peak. (**C**) Superposed MMN (deviant—standard) waveforms. Green shadow represents approximation of window taken for MMN analysis. (**D**) Bar plot of MMN responses (Naive-Keta z =-2.10, p =0.036; Naive-Veh z =0.420, p =0.674; Veh-Keta z =2.521, p =0.012; n=8) (**E**) Mean response to the standard before the deviant (SBD, dotted) and first standard (S1, solid) sound for naive (left), vehicle injected (middle), and ketamine injected (right) mice. (**F**) Bar plots show N1(Naive-Keta z =1.153, p=0.249; Naive-Veh z =-1.572, p=0.116; Veh-Keta z =-0.524, p=0.600; n=8) and P1(Ctrl-Keta z =-0.314, p=0.753; Naive-Veh z =0.105, p=0.917; Veh-Keta z =-0.105, p=0.917; n=8) amplitude following standard before the deviant (SBD) versus first (S1) sound stimulation in mice. (**G**) Bar plot of total distance travelled (Ctrl-Keta z =0.105, p =0.917; Naive-Veh z =0.734, p=0.463; Veh-Keta z =-0.105, p=0.917; n=6). Statistical comparisons were performed using the non-parametric Wilcoxon signed-rank test for paired measures.

**Figure 2.**
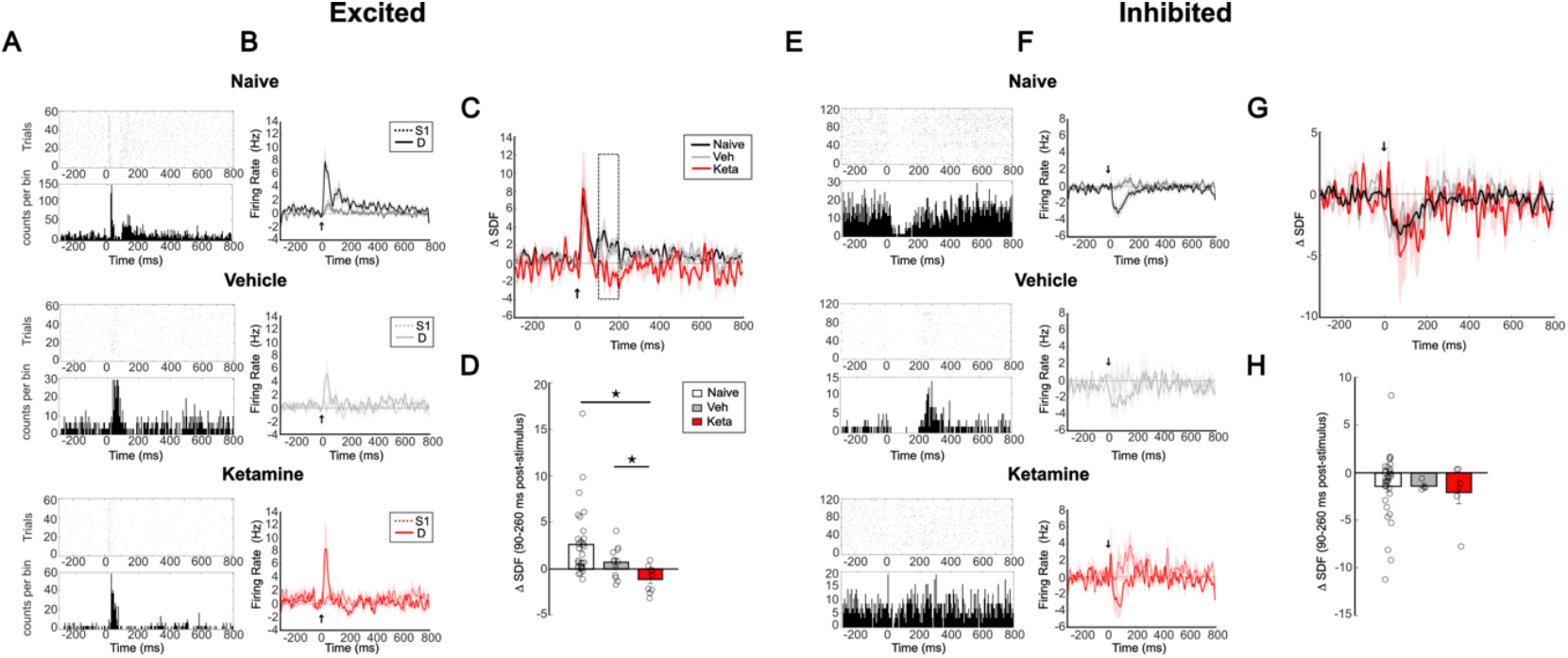
Subanesthetic ketamine alters spiking activity of responding neurons during deviant detection and stimulus specific adaptation. (**A, E**) Raster plots (top) and peristimulus time histograms (PSTH, bottom) for representative responding excited (**A**) and inhibited (**E**) neurons in naïve (top-black), vehicle injected (middle-gray), and ketamine injected (bottom-red) mice. (**B, F**) Spike density functions (SDF) of responding excited (**B**) and inhibited (**F**) neurons in each condition. (**C, G**) ΔSDF (D-S1) of responding excited (**C**) and responding inhibited cells (**G**) neurons in naive (top), vehicle injected (middle), and ketamine injected (bottom) mice. (**D, H**) Bar plots of evoked firing responses of responding excited cells calculated from ΔSDF during the time window 90-260 ms following stimuli presentation(**D**; Naive-Keta p= 0.034, mean difference= 2.950, std. error=1.346; Naive-Veh p= 0.029, mean difference= 1.549, std.error=0.658; Veh-Keta p=0.323, mean difference=-1.18, std.error=1.18) and responding inhibited cells (**H**; Naive-Keta p= 0.660, mean difference= 0.584, std. error=1.317; Naive-Veh p= 0.791, mean difference=0.383, std.error=1.432; Veh-Keta p=0.639, mean difference=0.674, std.error=1.384). A total of 5 mice were analyzed with the number of responding excited cells (naïve n=36, vehicle-injected n=12, ketamine-injected n=9) and responding inhibited cells (naïve n=33, vehicle-injected n=4, ketamine-injected n=6) varying among conditions. S1=first standard sound; D=deviant sound. Statistical comparisons were performed using Linear mixed model.

Conversely, neurons that were inhibited specifically by deviant sounds showed a slower and monophasic negative response that was not affected by ketamine administration (Figure 2E-H). These findings indicate that there are at least two different neuronal populations in A1 which differ in their sensitivity to ketamine. Moreover, they reveal that stimulus-specific inhibitory responses may persist despite reduced NMDA signaling, while novelty-specific excitatory responses require intact NMDA receptor function, suggesting a critical role for NMDA-mediated synaptic transmission in encoding auditory novelty.

### Oscillatory activity evoked by deviant sounds is significantly reduced by sub-anesthetic ketamine administration

Recordings from primary sensory areas across species shows that sensory stimulation reliably evokes a response in the gamma spectrum (centered at 40Hz), reflecting enhanced sensory processing during early perceptual encoding (Mulert *et al*., 2007; Naatanen *et al*., 2007; Polomac *et al*., 2015; Picard *et al*., 2019). To investigate whether ketamine administration affects this response in A1, we evaluated the oscillatory activity following standard and deviant sound presentations. Although the baseline oscillatory activity was similar between groups (Figure 3A-B), naive and vehicle-injected animals appeared to have a strong response in low gamma evoked specifically by deviant, but not standard, sounds (Figure 3C, left and middle panels). To facilitate comparing the response in the different experimental groups, we calculated the difference (Δ) in evoked PSD, (referred to as Δ PSD), by subtracting the following sound conditions: D-SBD (deviant detection) and S1-SBD (habituation) (Figure 3D, 3G). Δ-PSD showed a prominent narrow band peak at 40Hz in naive and vehicle injected animals, while ketamine administration was sufficient to significantly reduce this evoked response (Figure 3D, 3E). The effect of ketamine administration was similar when mice were exposed to the reversal oddball sequence (Supplemental Figure 3). Further, we did not observe any effects of ketamine on habituation (Figure 3F, G, H).

**Figure 3.**
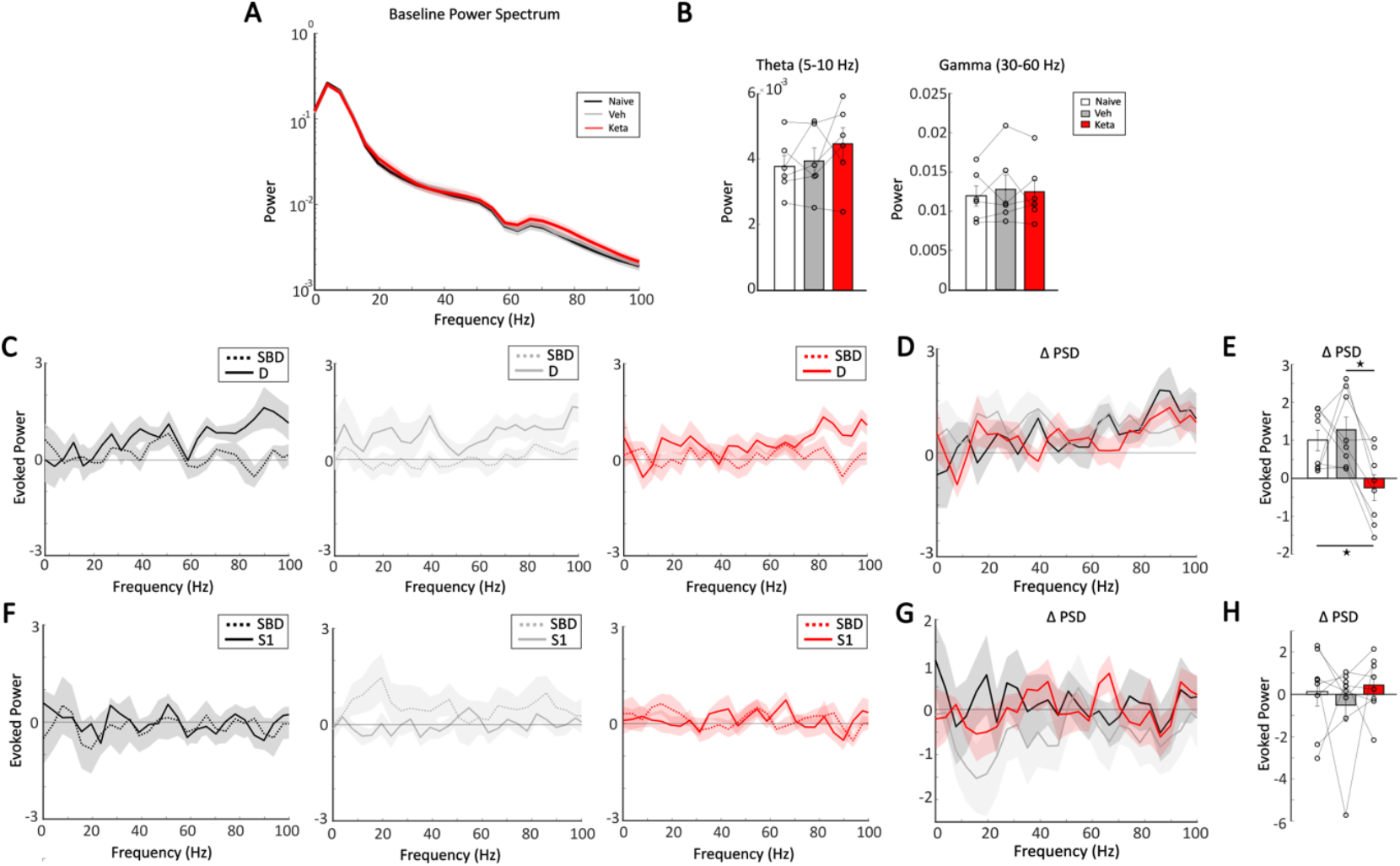
Subanesthetic ketamine affects power density spectra during deviance detection. (**A**) Power density spectra (0–100Hz) baseline (non-stimulus) EEG signal from the auditory cortex during immobility periods. A logarithmic scale is used for visualization purposes. (**B**) Bar plots show mean ± SEM power of theta (Naive-Keta z= 0.524, p=0.600; Veh-Keta z=-0.105, p=0.917; Ctrl-Veh z=0.943, p=0.345, n=6) and gamma (Naive-Keta z= 1.572, p=0.116; Veh-Keta z=0.943, p=0.345; Ctrl-Veh z=0.524, p=0.600, n=6) bands. (**C**) Baseline corrected evoked power in response to standard before the deviant (SBD) and deviant (D) sounds in naive (left, black), vehicle-injected (middle, gray), and ketamine-injected (right, red) mice. (**D**) ΔPSD (D-SBD) (**E**) Bar plot showing mean ± SEM ΔPSD at 40 Hz (Naive-Keta z= -2.240, p=0.025; Veh-Keta z=2.380, p=0.017; Naive-Veh z=0.140, p=0.889; n=8). (**F**) Evoked power in response to the standard before the deviant (SBD) and first standard (S1) sounds in naive (left, black), vehicle-injected (middle, grey), and ketamine-injected (right, red) mice. (**G**) ΔPSD (SBD-S1). (**H**) Bar plot showing mean ± SEM ΔPSD at 40 Hz (Naive-Keta z= -2.521, p=0.012; Veh-Keta z=1.820, p=0.069; Naïve-Veh z=1.680, p=0.093; n=8). Statistical comparisons were performed using the non-parametric Wilcoxon signed-rank test for paired measures.

Therefore, altogether, these results suggest that sub-anesthetic ketamine generally disrupts the brain’s ability to respond to unpredictable stimuli in auditory input streams.

### Sub-anesthetic ketamine administration disrupts A1-PPC functional connectivity elicited by auditory deviance detection

Emerging evidence suggests that the PPC, a higher-order associative region involved in both multisensory integration and sensorimotor coordination (Andersen & Buneo, 2002; Van Derveer *et al*., 2023), is implicated in top-down modulation of sound predictability (Hockley *et al*., 2025). Therefore, we next investigated whether PPC integrates rule violation in the auditory stream. To do so, we analyzed MMN waveforms (D-SBD) and observed a prominent MMN response in naïve mice, similar to A1. Ketamine (Figure 4A-C), but not vehicle (Figure 4D-F), administration induced a significant reduction of the MMN response in the PPC, again concurrent to findings in A1. The effect of ketamine administration was similar when mice were exposed to the reversal oddball sequence (Supplemental Figure 4). These results indicate that PPC integrates rule-violation stimuli, and its responses to deviant sounds require intact NMDA receptor-mediated signaling. Furthermore, neither ketamine nor vehicle administration had any significant influence on auditory habituation (Figure 4G-J), suggesting that the PPC is not involved in habituation responses to repetitive stimuli.

**Figure 4.**
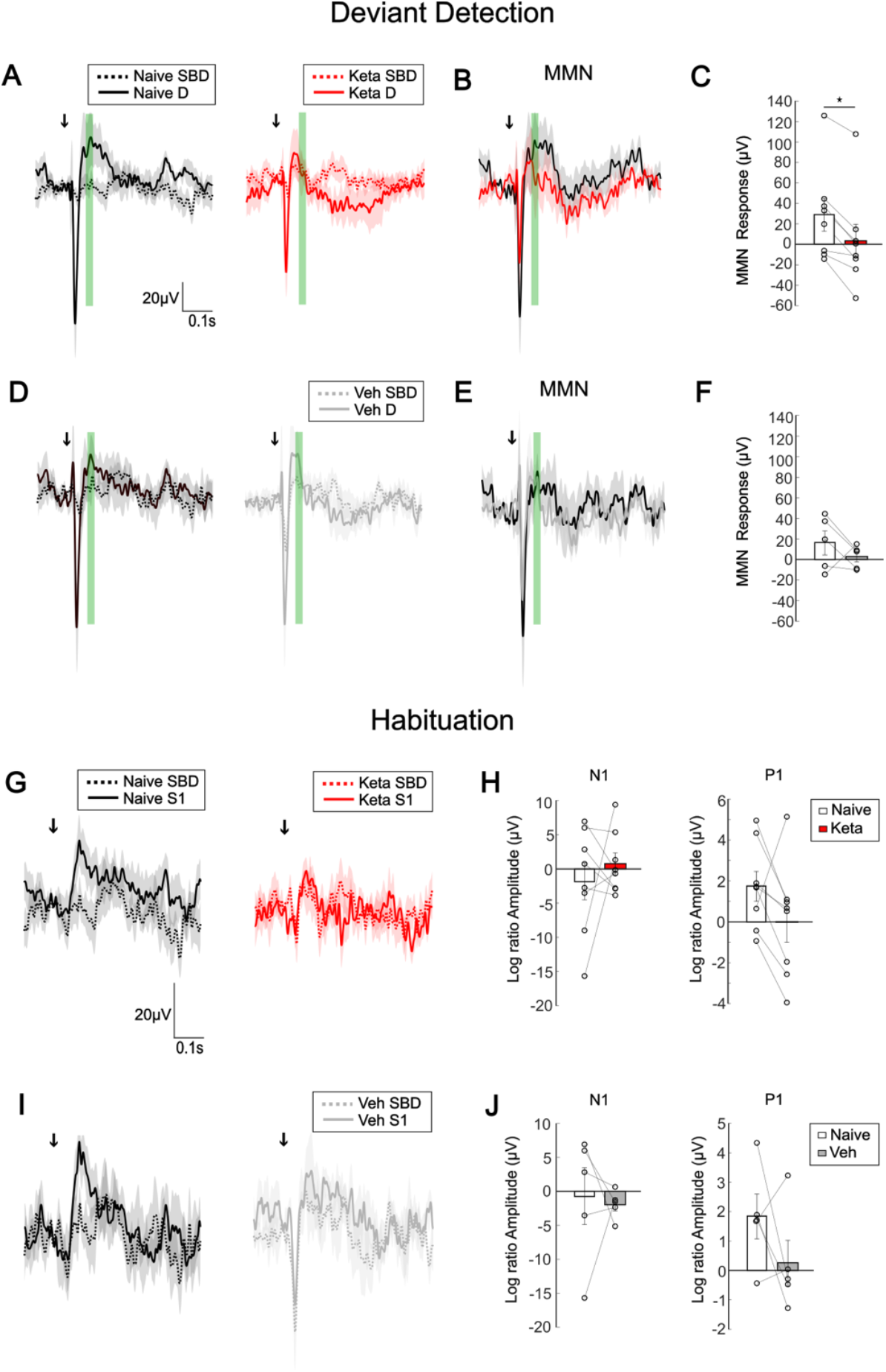
Subanesthetic ketamine reduces deviant detection encoding in the PPC. (**A, B**) Deviant detection in the PPC of naive versus ketamine-injected mice during oddball protocol. (**A**) Mean ERP traces of standard before the deviant (SBD, dotted) and deviant (D, solid) sounds for naive (left) versus ketamine injected (right) mice. Green shadow represents time window taken for MMN analysis. (**B**) Superposed MMN (deviant - standard) waveforms. (**C**) Bar plot of MMN responses (Wilcoxon signed ranked test: z=-2.521, p=0.012; n=8) in naïve (white) and ketamine (red) -injected mice. (**D**-**F**) Deviant detection in the PPC of naive vs vehicle-injected mice. (**D**) Mean ERP traces of standard before the deviant (SBD, dotted) and deviant (D, solid) sounds for naive (left) and vehicle injected (right) mice. (**E**) Superposed MMN (deviant—standard) waveforms. (**F**) Bar plot of MMN responses (Wilcoxon signed ranked test: z=-1.214, p=0.225; n=5) in naïve (white) and vehicle -injected (grey) mice. (**G**) Mean ERP traces of the standard before the deviant (SBD, dotted) and first standard (S1, solid) sounds for naïve (left) versus ketamine-injected (right) mice. (**H**) Bar plots of S1/SBD log ratio N1(Wilcoxon signed ranked test: z=-0.135, p=0.893; n=8) and P1 (Wilcoxon signed ranked test: z=0.674, p=0.500; n=8) amplitude in naive and ketamine-injected mice. (**I**) Mean ERP traces of the standard before the deviant (SBD, dotted) and first standard (S1, solid) sounds for naïve (left) versus vehicle injected (right) mice. (J) Bar plots of S1/SBD log ratio N1(Wilcoxon signed ranked test: z=1.069, p=0.285; n=5) and P1 (Wilcoxon signed ranked test: z=1.069, p=0.285; n=5) amplitude in naive and vehicle-injected mice.

To explore the effects of ketamine on inter-cortical communication during deviance detection, we performed weighted Phase Lag Index (wPLI) analyses during the presentation of deviant and standard sounds. wPLI assesses the lagged phase synchronization between signals, offering a robust measure of functional connectivity (Vinck *et al*., 2011). We applied this method to the time window corresponding to the MMN response (80-320ms after stimuli presentation), a time window chosen to avoid erratic or sudden changes in the signal due to the non-stationary nature of the evoked potentials.

wPLI analysis showed an increased functional connectivity between A1 and PPC specifically following deviant presentation, in both naïve and vehicle injected mice (Figure 5). This increased connectivity was specifically at 30-38 Hz for the oddball sequence consisting of 5kHz standard tone and 10kHz deviant tone (Figure 5B, C), while it was at 50-55 kHz for the reversal oddball sequence (10kHz standard tone and 5 kHz deviant tone; Figure 5 E, F). This increased connectivity between A1 and PPC was abolished following ketamine administration (Figure B, E). These results suggest an important role of A1-PPC connectivity underlying deviance detection but not habituation.

**Figure 5.**
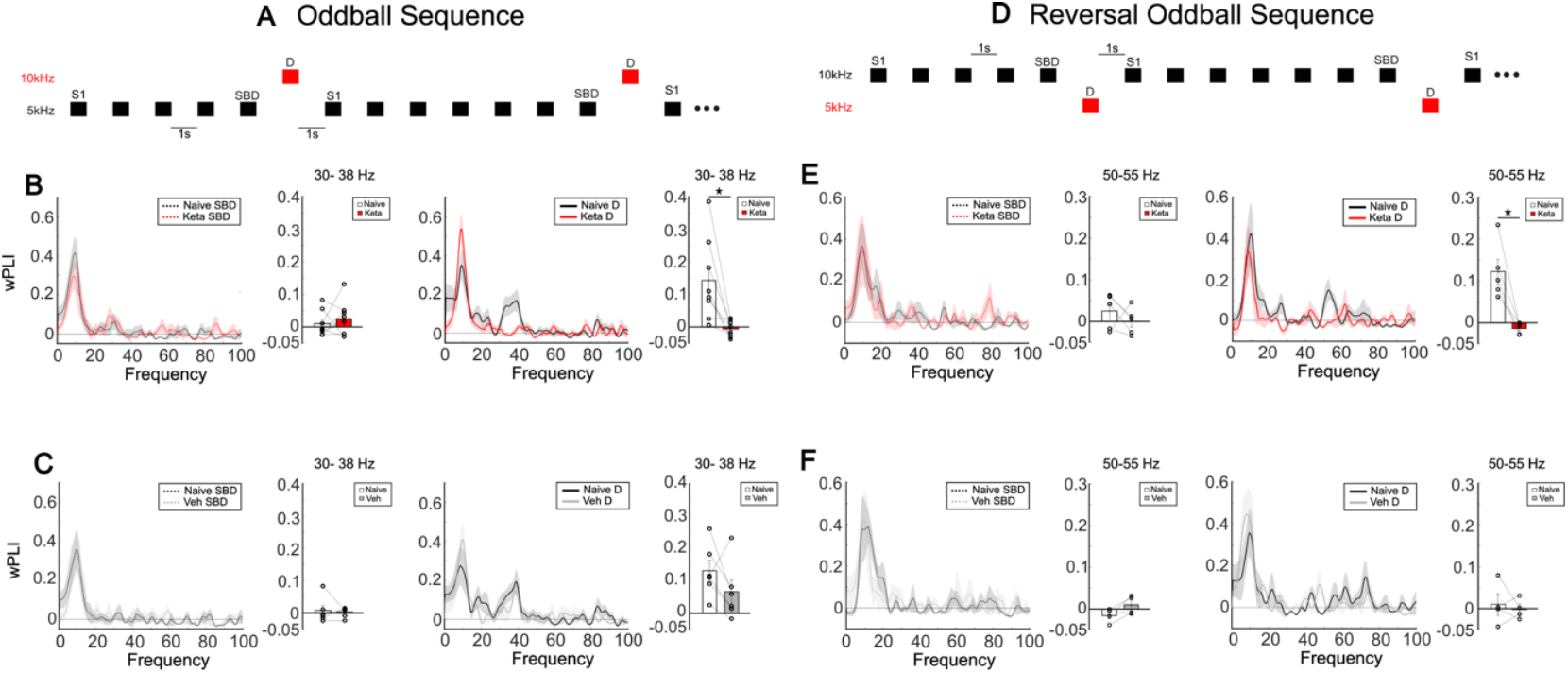
Sub-anesthetic ketamine administration disrupts A1-PPC functional connectivity elicited by auditory deviance detection. wPLI between A1 and PPC during oddball protocol. (**A**) Schematic of oddball protocol used in (B-C). (**B**) wPLI spectrum between A1 and PPC and bar plot showing wPLI at 30-38Hz for the standard before the deviant (SBD) sound (left) (z=0.700, p=0.484; n=8) and deviant sound (D, right) (z= -2.380, p=0.017; n=8) for naive versus ketamine conditions. (**C**) wPLI spectrum between A1 and PPC and bar plot showing wPLI at 30-38Hz for the standard before the deviant sound (SBD, left) (z=-1.153, p=0.249; n=6) and deviant sound (D, right) (z=-1.153, p=0.249; n=6) for naive versus vehicle conditions. **(D-F)** wPLI spectrum between A1 and PPC during reversal oddball sequence. (**D**) Schematic of reversal oddball protocol used in (E-F). (**E**) wPLI spectrum between A1 and PPC and bar plot showing wPLI at 50-55Hz for the standard before the deviant sound (SBD, left) (z=-0.674, p=0.500; n=8) and deviant sound (D, right) (z=-2.023, p=0.043; n=8) for naive versus ketamine conditions. (**F**) wPLI spectrum between A1 and PPC and bar plot showing wPLI at 50-55Hz for the standard before the deviant (SBD) sound (left) (z=1.461, p=0.144; n=6) and deviant sound (right) (z=0.000, p=1.000; n=6) for naive versus vehicle conditions. SBD=standard sounds before the deviant; D=deviant sound. Statistical comparisons were performed using the non-parametric Wilcoxon signed-rank test for paired measures.

Overall, this data suggests that rule violation in the auditory stream elicits functional communication between A1 and PPC, which is dependent on intact NMDA-mediated signaling.

## DISCUSSION

The core concept of predictive coding theory suggests that the brain generates a model of the environment based on its previous experiences and forms a predictive coding framework. Incoming sensory information is continuously referenced to this internal predictive model, and cases of mismatch give rise to a prediction error signal. The auditory oddball paradigm (repeating standard sounds randomly interrupted by rare deviant stimuli) has been routinely used to elucidate how the brain encodes regular auditory patterns, generates inherent models of predictability and uses it to efficiently detect deviations from these internal models (Naatanen *et al*., 2007; Fitzgerald & Todd, 2020).

MMN, the electrophysiological response to rule violations, is a well-established biomarker of auditory deviant detection, whereby an auditory evoked potential to a deviant stimulus is greater than the response to repetitive standard stimuli. Using this biomarker, we found that sub-anesthetic ketamine administration significantly reduced the amplitude of the auditory MMN response. In addition, oscillatory analyses further revealed that ketamine markedly reduced the narrow low gamma response selectively evoked by deviant sounds, impairing novelty detection at the gamma-band level. These findings validate previous work demonstrating that MMN depends on intact NMDA receptor signaling across species (Umbricht *et al*., 2000; Gil-da-Costa *et al*., 2013; Chen *et al*., 2015; Javitt & Sweet, 2015; Rosburg & Kreitschmann-Andermahr, 2016; Schuelert *et al*., 2018). Nevertheless, the underlying neural and inter-cortical network mechanisms are still under debate. To bridge this gap, our study extends beyond classical population-level measures by uncovering the cellular correlates of deviance detection and the mesoscale connectivity that links primary sensory and associative cortices.

Our results at the cellular level show that the firing response of a subpopulation of A1 neurons follows a biphasic pattern: a rapid initial peak shortly after stimulus presentation (0-95 ms after stimulus onset), followed by a slower later peak (between 100 ms and 200 ms), that coincides with the timing of the MMN response at the population level. This biphasic pattern consistently appeared during deviant sounds, while standard sounds only elicited fast monophasic responses. Furthermore, sub-anesthetic ketamine administration significantly and specifically reduced the second peak of spiking activity following the presentation of a deviant sound. These data suggest that ketamine administration has a selective effect on the second slower peak of the response, presumably reflecting a higher-order realization of sound processing. It is possible that slower, ketamine-sensitive peak of neuronal firing could be due to disinhibition of cortical excitatory activity. A greater NMDA sensitivity of inhibitory circuits could then explain why this second peak is abolished following sub-anesthetic ketamine administration (Grunze *et al*., 1996; Li *et al*., 2002; Homayoun & Moghaddam, 2007; Middleton *et al*., 2008; Widman & McMahon, 2018; Picard *et al*., 2019). For example, previous studies have shown that sub-anesthetic ketamine administration preferentially inhibits PV+ interneurons, by targeting NMDA receptors containing the 2A subunit, and regulates evoked gamma-band oscillations in visual cortex (Picard *et al*., 2019). This inhibitory effect of ketamine on PV+ interneurons could indeed explain our observations of a reduced evoked 40Hz response to deviant sounds, since reduced evoked 40Hz rhythms have been linked to PV+ interneuron hypofunction (Cardin *et al*., 2009; Picard *et al*., 2019; Batista-Brito *et al*., 2023). Additionally, long-range callosal PV+ neurons have been shown to play a critical role in inter-area connectivity, in different cortical regions, including the prefrontal (Cho *et al*., 2023) and auditory cortex (Zurita *et al*., 2018). Other GABAergic interneurons populations, including VIP and SST+ interneurons have been also involved in disinhibitory mechanisms mediated by neuromodulators (reviewed in(Kullander & Topolnik, 2021); however, whether and how the activity of different interneuron populations is affected by acute sub-anesthetic ketamine treatment remains to be elucidated. Interestingly, sub-anesthetic ketamine administration can bidirectionally modulate cortical acetylcholine levels (Kikuchi *et al*., 1997; Kim *et al*., 1999). Cholinergic, and in particular, nicotinic stimulation has been demonstrated to enhance MMN amplitudes (Harkrider & Hedrick, 2005; Inami *et al*., 2005; Inami *et al*., 2007; Martin *et al*., 2009; Hamilton *et al*., 2018; Weber *et al*., 2022). Therefore, altered basal forebrain to A1 communication, possibly through cross-talk with dis-inhibitory microcircuits could contribute to the specific effect of ketamine on the slower later peak of spiking activity.

The second peak of A1 neuron firing elicited by the deviant sound could be also mediated by astrocytic signaling. For example, in the lateral habenula, foot-shock stress has been shown to induce a bimodal neuronal response, similarly to what we observed in A1 neurons following deviant sounds (Xin *et al*., 2025). In this region, the second peak of neuronal activation was shown to be mediated by astrocytic release of the gliotransmitters glutamate and ATP/adenosine (Xin *et al*., 2025). Further, a recent study has shown that neuromodulators can signal through astrocytes to modulate synapse strength in adult mouse hippocampus (Lefton *et al*., 2025). Whether astrocytic signaling contribute to MMN responses in A1 is thus far unknown.

Recent studies emphasize the importance of whole-brain connectivity for processing auditory stimuli. While the frontal cortex has been extensively studied for its role in top-down control of prediction errors (Hockley *et al*., 2025), far less is known about the PPC. As a hub for multisensory integration (Choi & Lee, 2025) and higher-order auditory processing (Andersen & Buneo, 2002; Van Derveer *et al*., 2023), the PPC has been shown to respond to unexpected sounds (Justen & Herbert, 2018; Mohan *et al*., 2018; Raltschev *et al*., 2025); however, its precise role in differentiating redundant from deviant auditory stimuli remains largely unexplored. In this study, we show that PPC exhibited a strong response to sounds. As it was the case in A1, in PPC ketamine administration significantly decreased the amplitude of MMN, which is thought to carry the novelty content of the response. Based on this observation, we focused our connectivity analyses on the late response period. Our wPLI analysis revealed a consistent low-frequency peak around 8–10 Hz in A1–PPC connectivity, which was present across conditions and is likely to reflect a baseline, task-independent coupling between auditory and parietal cortices, as previously reported in resting-state EEG studies (Wang *et al*., 2017). In contrast, an additional coupling peak emerged selectively in the gamma range during deviant (unexpected) sounds, with its frequency depending on deviant stimulus identity (≈30 Hz for 10 kHz and ≈50 Hz for 5 kHz tones), and was absent when the same sounds were presented as repetitive standards. We interpret this stimulus-locked, frequency-specific narrow gamma synchronization as reflecting a transient, expectation-violation signal carrying information about the stimulus probability (“this sound was not expected”), while the precise gamma band engaged may encode stimulus identity (“this sound is 5 kHz vs. 10 kHz”). This effect may arise because ketamine administration abolishes the functional connectivity between A1 and PPC, by altering either A1 local circuits primarily and/or communication between A1 and PPC. These observations provide, to the best of our knowledge, the first evidence that functional connectivity between A1 and PPC supports auditory deviance detection, highlighting a previously uncharacterized mesoscale pathway for rule-deviation processing.

Overall, our observations are consistent with previous studies reporting decreased directed functional connectivity across the brain under various psychedelics drugs (Muthukumaraswamy *et al*., 2015; Barnett *et al*., 2020). These data further support the hypothesis that the psychedelic state involves a breakdown in patterns of functional organization or information flow in the brain (Barnett *et al*., 2020). Paradoxically, ketamine’s ability to restore impaired cortical networks, particularly in regions such as the prefrontal cortex, is considered central to its antidepressant effects in treatment-resistant depression (Li *et al*., 2016; Roussy *et al*., 2021), (Abdallah *et al*., 2017; Gass *et al*., 2019). This may be due to the specific nature of the underlying pathophysiology of treatment-resistant depression. Intriguingly, MMN has been shown to be impaired in psychiatric disorders such as depression (Tseng *et al*., 2021) and schizophrenia (Molnar *et al*., 2024), both of which have been linked to NMDA receptor dysfunction, although in opposite directions. In schizophrenia, NMDA receptor hypofunction is thought to contribute to reduced MMN amplitude and sensory processing deficits, while in depression, NMDA receptor hyperfunction is believed to underlie similar sensory processing disruptions (Balu, 2016; Adell, 2020; Wang *et al*., 2022). Although, the therapeutic mechanisms behind ketamine’s anti-depressive effects are still not well understood, they are thought to involve multiple parallel mechanisms, including altered neurotransmission through NMDA receptor antagonism, enhanced synaptic plasticity and modulation of various signaling pathways, such as the mTOR pathway (Duman *et al*., 2016).

In summary, our data identify a specific biphasic spiking response in a subpopulation of A1 neurons elicited by deviant, but not standard sounds, with the second peak dependent on NMDA receptor–mediated signaling. The identity of the neurons participating in this response and the underlying microcircuit mechanisms remain to be elucidated. We further provide evidence that the PPC responds to deviant, but not repetitive, sounds and that this response is dependent on NMDA receptors. Finally, we demonstrate a functional connectivity between A1 and PPC likely contributing to the integration of deviant sounds, which is abolished by sub-anesthetic ketamine administration. Together, these data support the hypothesis that PPC, through intact NMDA receptor signaling, contributes to the mesoscale network underlying prediction error processing. By bridging cellular, population, and mesoscale levels, our study offers the first integrated evidence of how NMDA-dependent auditory circuits shape rule-deviation processing, unveiling a previously uncharacterized pathway for novelty detection. A caveat of our study is that IP administration of ketamine has a system-wide effect, precluding the identification of specific circuit and cellular location of NMDA receptors involved in deviance detection; this open question should be addressed by future experiments. Further studies are also needed to understand the precise role of A1-PPC connectivity and PPC itself in the processing of rule-violation processing.

## ACKNOWLEDGEMENTS

We would like to thank Kristian Agbogba, Mikael Valton Charette and James Waldron for their technical assistance. We would also like to thank the Comité Institutionnel de Bonne Pratiques Animales en Recherche (CIBPAR), all the personnel of the animal facility and the Plateforme Imagerie Microscopique (PIM) of the Azrieli Research Center of CHU Sainte-Justine, and Compute Canada for their instrumental technical support and members of our lab for insightful data discussion. We are grateful to Dr Jozsef Csicsvari and his lab for sharing their spike sorting analysis pipeline. This work was supported by La Fondation des Étoiles (G.DC.); the Canadian Neurodevelopmental Research Training (CanNRT) Platform (M.I.C-M and A.C.C) and the Savoy Foundation fellowship (M.I.C-M).

## DATA AVAILABILITY

All data and codes are available upon request.

## AUTHORS CONTRIBUTION

MIC-M, BC, GDC designed the experiments. MIC-M, ACC, PC performed the experiments. MIC-M, ACC analyzed data. MIC-M, ACC, BC, GDC wrote the manuscript. All authors read and corrected the manuscript.

