## Supplemental Data for "Sub-anesthetic ketamine administration decreases deviance detection responses at the cellular, populational and mesoscale levels"

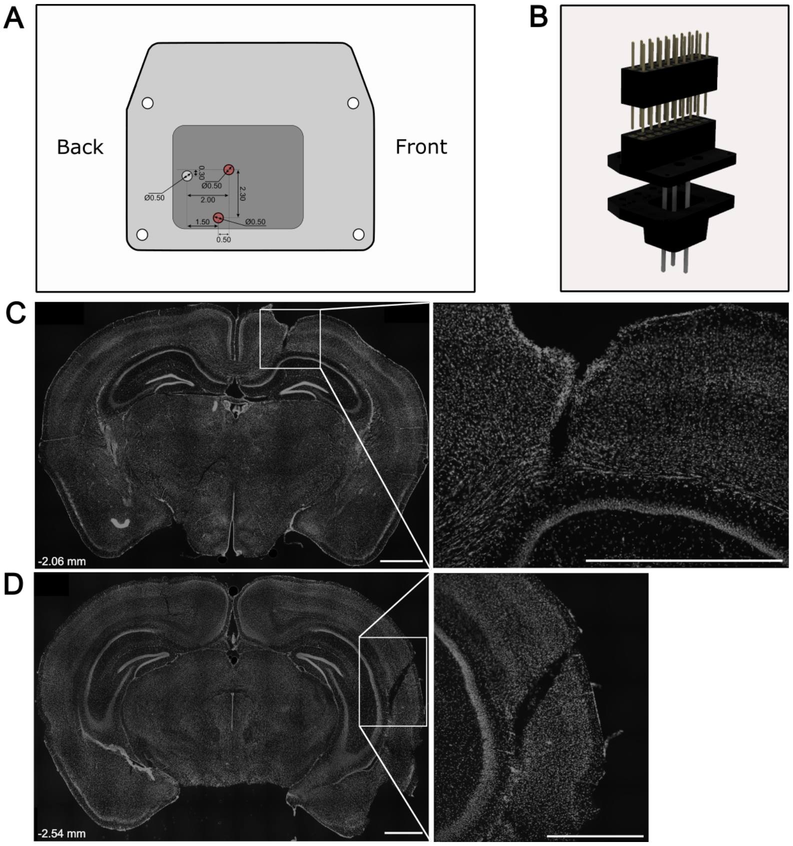


**Supplemental Figure 1. Microdrive geometry and electrode bundle location**. **A)** Schematic representation of electrode cluster positions within the custom-made 3D-printed microdrive-like scaffold. Arrows indicate the distances between hole positions. Electrode clusters implanted for recordings in the posterior parietal cortex (PPC) and primary auditory cortex (A1) are shown in red. **(B)** Schematic showing a 3D representation of the microdrive-like structure and the milmax connector used in this study. **C**, **D**) Whole brain section (left) and high magnification (right) images showing electrode placement for PPC **(C)** and A1 **(D)**. Cell nuclei are labelled by DAPI (gray). Coordinates from bregma are indicated on the lower left corner. Scale bars: 1mm.

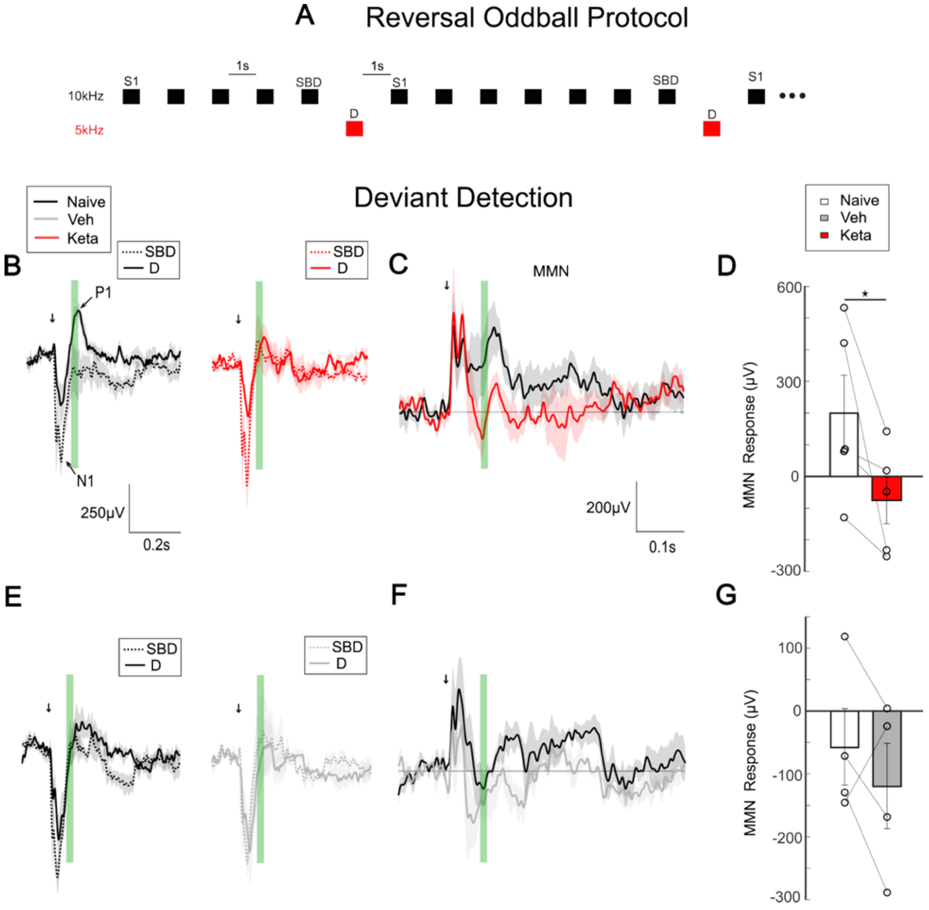


**Supplemental Figure 2. Subanesthetic ketamine alters MMN responses during deviant detection in the reversal oddball sequence.** (**A**) Schematic representation of the reversal oddball protocol with black squares representing 10kHz pure tones (standard sounds) and the red squares 5kHz pure tones (deviant sounds). (**B**) Mean evoked related potentials (ERPs) traces of standard (dotted) and deviant (solid) sounds for naive (left) and ketamine injected (right) mice. Arrows indicate the start time of stimulation. N1 is identified as the first negative peak and P1 as the first positive peak. (**C**) Superposed MMN (deviant—standard) waveforms. (**D**) Bar plot of MMN response of naïve versus ketamine injected mice (Wilcoxon signed ranked test; z =-2.023, p =0.043; n=5) (**E**) Mean ERP traces of standard (dotted) and deviant (solid) sounds for naive (left) and vehicle injected (right) mice. (**F**) Superposed MMN (deviant—standard) waveforms. (**G**) Bar plot of MMN response of naïve versus vehicle injected mice (Wilcoxon signed ranked test; z =-0.720, p =0.465; n=4). Green shadow represents window taken for MMN analysis (see methods). SBD=standard sounds before the deviant; D=deviant sound.

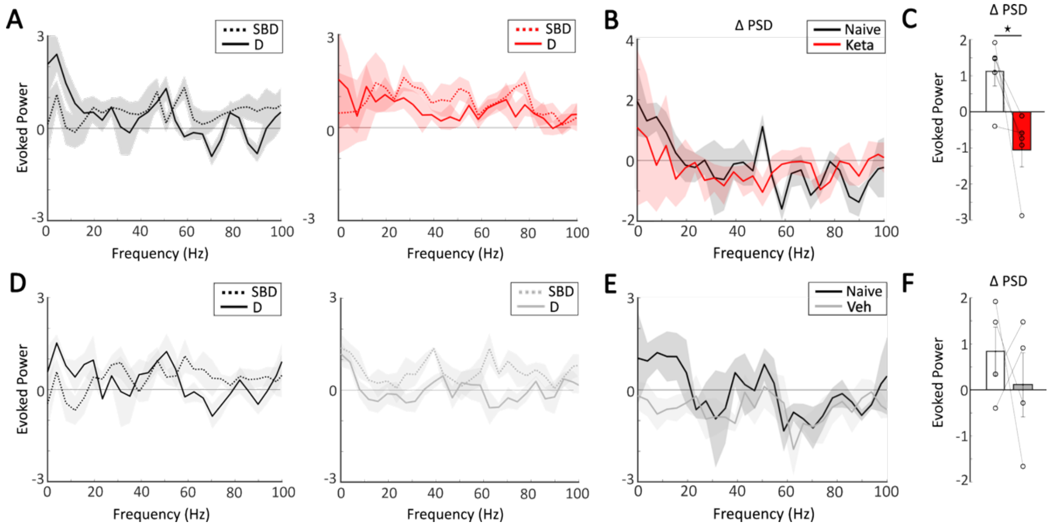


**Supplemental Figure 3. Subanesthetic ketamine affects power density spectra during deviance detection in the reversal oddball sequence.** (**A**) Evoked power in response to standard (SBD) and deviant (D) sounds in naive (left) and ketamine-injected (right) mice. (**B**) ΔPSD of evoked power in response to standard and deviant sounds across all frequencies for naïve versus ketamine injected mice. (**C**) Bar plot of ΔPSD of evoked power at 50 Hz for naïve versus ketamine-injected mice (Wilcoxon signed ranked test: z= -2.023, p=0.043, n=5). (**D**) Evoked power in response to standard (SBD) and deviant (D) sounds in naive (left) and vehicle-injected (right) mice. (**E**) ΔPSD of evoked power in response to standard and deviant sounds across all frequencies for naïve versus vehicle injected mice. (**F**) Bar plot of ΔPSD of evoked power at 50 Hz for naïve versus vehicle-injected mice (Wilcoxon signed ranked test: z= -0.730, p=0.465, n=4).

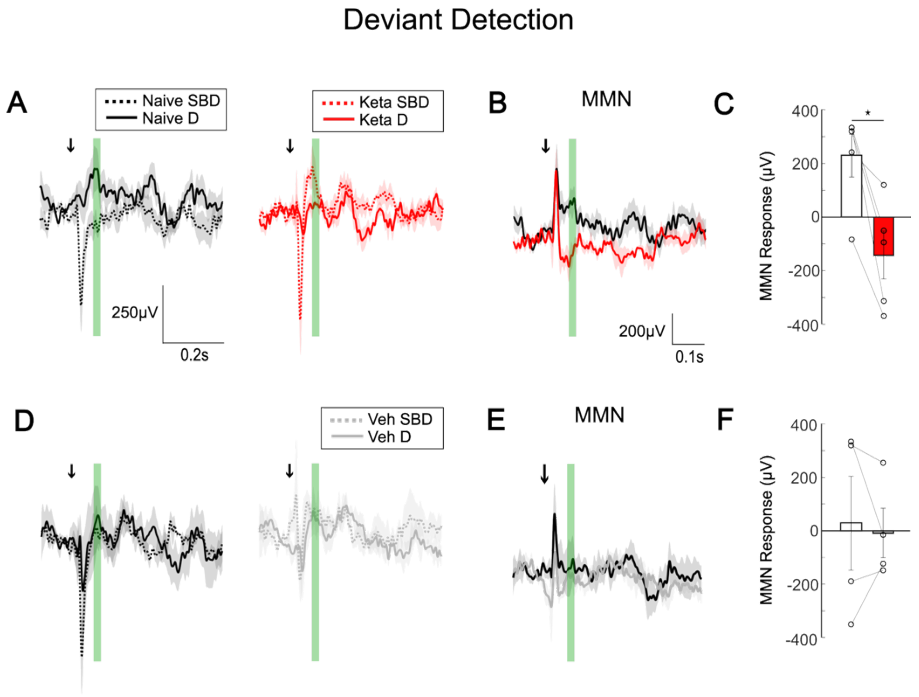


**Supplemental Figure 4. Subanesthetic ketamine reduces deviant detection encoding in the PPC in the reversal oddball sequence.** (**A**, **B**) Deviant detection in the PPC of naive versus ketamine-injected mice during reversal oddball protocol. (**A**) Mean ERP traces of standard (dotted) and deviant (solid) sounds for naive (left) versus ketamine injected (right) mice. (**B**) Superposed MMN (deviant—standard) waveforms. (**C**) Bar plot of MMN responses (Wilcoxon signed ranked test: z=-2.023, p=0.043; n=5) in naive and ketamine-injected mice. (**D**-**F**) Deviant detection in the PPC of naive vs vehicle-injected mice during reversal oddball protocol. (**D**) Mean ERP traces of standard (dotted) and deviant (solid) sounds for naive (left) and vehicle injected (right) mice. (**E**) Superposed MMN (deviant—standard) waveforms. (**F**) Bar plot of MMN responses (Wilcoxon signed ranked test: z=-0.365, p=0.715; n=4) in naive and vehicle-injected mice. Green shadow represents time window taken for MMN analysis. SBD=standard sounds before the deviant; D=deviant sound.
